# Identification of split-GAL4 drivers and enhancers that allow regional cell type manipulations of the *Drosophila melanogaster* intestine

**DOI:** 10.1101/2020.08.24.264887

**Authors:** Ishara S. Ariyapala, Jessica M. Holsopple, Ellen M. Popodi, Dalton G. Hartwick, Lily Kahsai, Kevin R. Cook, Nicholas S. Sokol

## Abstract

The Drosophila adult midgut is a model epithelial tissue composed of a few major cell types with distinct regional identities. One of the limitations to its analysis is the lack of tools to manipulate gene expression based on these regional identities. To overcome this obstacle, we applied the intersectional split-GAL4 system to the adult midgut and report 653 driver combinations that label cells by region and cell type. We first identified 424 split-GAL4 drivers with midgut expression from over 7,300 drivers screened, and then evaluated the expression patterns of each of these 424 when paired with three reference drivers that report activity specifically in progenitor cells, enteroendocrine cells, or enterocytes. We also evaluated a subset of the drivers expressed in progenitor cells for expression in enteroblasts using another reference driver. We show that driver combinations can define novel cell populations by identifying a driver that marks a distinct subset of enteroendocrine cells expressing genes usually associated with progenitor cells. The regional cell type patterns associated with the entire set of driver combinations are documented in a freely available website, providing information for the design of thousands of additional driver combinations to experimentally manipulate small subsets of intestinal cells. In addition, we show that intestinal enhancers identified with the split-GAL4 system can confer equivalent expression patterns on other transgenic reporters. Altogether, the resource reported here will enable more precisely targeted gene expression for studying intestinal processes, epithelial cell functions, and diseases affecting self-renewing tissues.

## INTRODUCTION

The Drosophila midgut is a model system for understanding tissue homeostasis and adaptation in response to environmental challenges, including dietary changes, alterations in microbiota, and the ingestion of pathogens and harmful chemicals (Miguel-Aliaga et al., 2018). While physiologically and functionally orthologous to the mammalian intestine, the fly midgut is comparatively simple and composed of only four main cell types (Zwick et al., 2019): intestinal stem cells (ISCs), their undifferentiated enteroblast (EB) daughters, and two differentiated cell types, absorptive enterocytes (ECs) and hormone-producing enteroendocrine cells (EEs) (Micchelli and Perrimon, 2006; Ohlstein and Spradling, 2006). These four cell types display distinct regional identities along the length of the midgut, which has been divided into five main regions (R1–R5) that carry out the sequential processes of food breakdown, nutrient absorption, and waste elimination (Buchon et al., 2013; Marianes and Spradling, 2013). In contrast to most tissues, which show little cell turnover during adult life, the midgut is continually remodeling through careful adjustments to stem cell division rates, division patterns, and differentiation decisions. These dynamic cell composition changes make the midgut exceptionally well suited for studying cellular processes common to all self-renewing tissues under normal and pathological states.

A current limitation to the use of the Drosophila midgut as a model system is the lack of tools to manipulate gene expression in a cell type and/or regional manner. The most common tool for experimentally controlling gene expression in Drosophila is the GAL4–UAS system (Brand and Perrimon, 1993). Pioneering studies reported the intestinal expression profiles of dozens of GAL4 drivers and identified several that were expressed in specific regions (Buchon et al., 2013; Marianes and Spradling, 2013), but many of these drivers have limited utility because their cell type expression was not characterized or they express in multiple cell types. In addition, few midgut enhancer sequences have been molecularly characterized, so it is difficult to design new transgenes targeting defined cells. Most known enhancers do not limit expression to a specific cell type (*e*.*g*., the *mex1* enhancer, present in a widely used EC driver, also directs expression in stem cells and EBs (Lucchetta and Ohlstein, 2017)), or they do not direct expression in all cells of a single type (*e*.*g*., EE enhancers from *AstA, AstC, Dh31* and *Tk* and EC enhancers from *Myo31DF* drive expression in only a subset of these cell types (Beehler-Evans and Micchelli, 2015; Chen et al., 2016)). Better tools would enable investigators to target specific cells for genetic experimentation based on their type and their position within biochemically and physiologically specialized intestinal regions.

We have addressed this limitation by developing a strategy for characterizing the intestinal cell type expression profiles of split-GAL4 drivers and applying it to the large collection of split-GAL4 driver stocks maintained at the Bloomington Drosophila Stock Center (Dionne et al., 2018; Tirian and Dickson, 2017). In the split-GAL4 system, the DNA-binding domain of GAL4 and a transcriptional activator are expressed from separate transgenes with different regulatory sequences (Luan et al., 2006). Combining the two transgenes with simple crosses activates *UAS* transgenes only in cells where both regulatory sequences are active (Fig 1A). While regulatory sequences typically direct transcription in widespread and complex patterns, this split-GAL4 intersectional scheme often defines very limited sets of cells or even single cells. The expression of these split-GAL4 drivers has been characterized most extensively in the adult brain—a complex, terminally differentiated tissue with little cell turnover in adults (Chen et al., 2019; Dionne et al., 2018; Dolan et al., 2019; Sekiguchi et al., 2020; Wolff and Rubin, 2018). In this study, we extend the analysis of these drivers to the intestine. We first identify the subset of split-GAL4 drivers that are expressed in the midgut, and then characterize the cell type expression patterns of these drivers relative to a set of cell type-specific reference strains. We also show that enhancer sequences characterized by this approach can generate equivalent expression patterns when used in other transgenes. Altogether, this analysis identifies a large set of useful drivers and enhancer sequences that will enable more precise gene manipulations for studying intestinal processes, epithelial cell functions, and diseases affecting self-renewing tissues.

**Figure 1.**
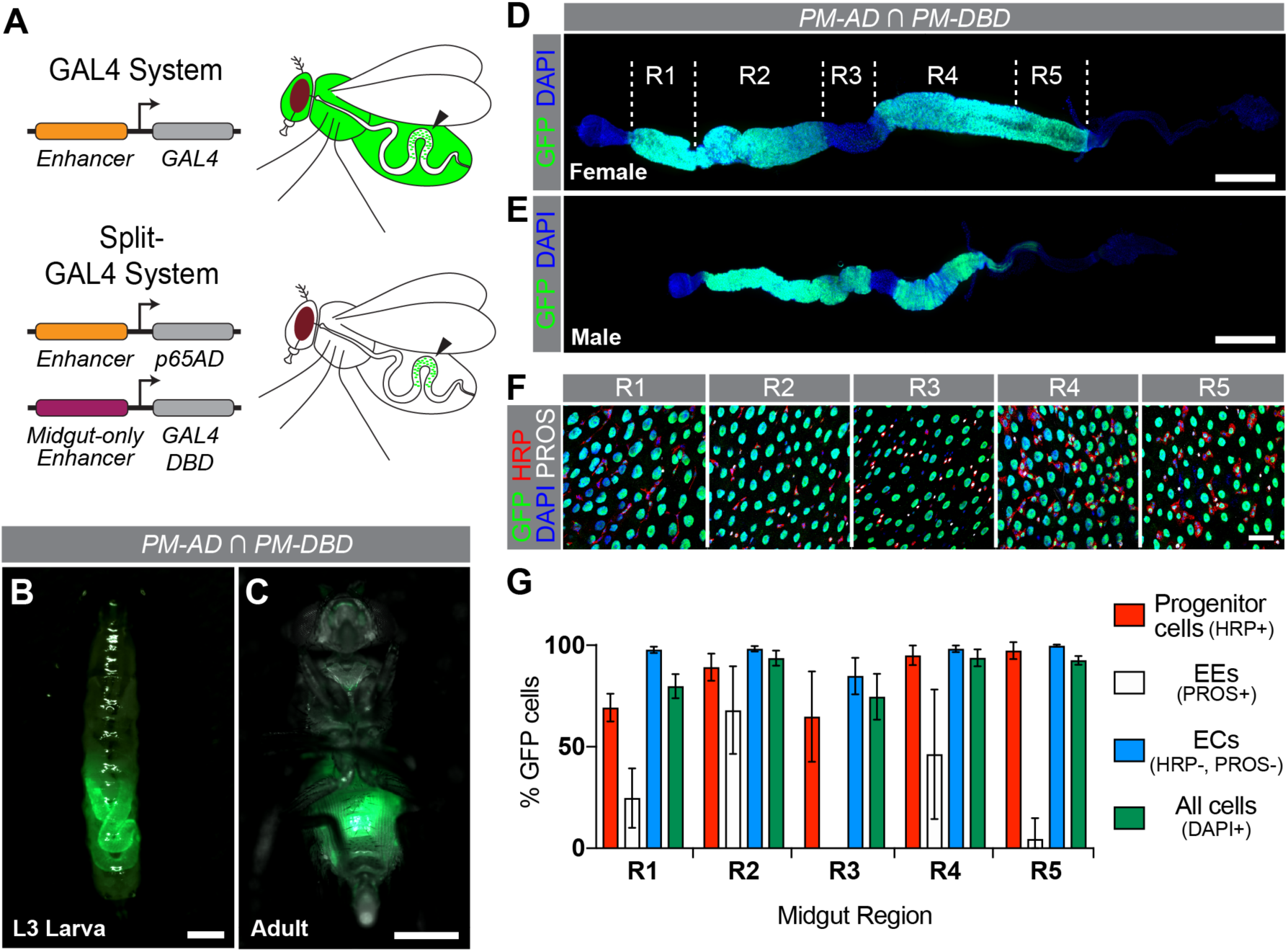
Pan-midgut split-GAL4 drivers label most cells in the Drosophila intestinal epithelium. (A) Identifying intestinally active enhancers using the split-GAL4 system. In this example, a midgut-only DBD reference driver restricts expression of *UAS-GFP* to a specific intestinal region when combined with a p65AD driver expressed more widely. (B, C) Larval and adult expression of *UAS-6XGFP* driven by pan-midgut (PM) split-GAL4DBD and p65AD drivers (*CG10116-p65AD* ∩ *CG10116-GAL4DBD*). (D, E) *UAS-6XGFP* expression driven by *PM-AD* and *PM-DBD* drivers in female (D) and male (E) intestines counterstained with DAPI (blue). (F) *UAS-GFP.nls* expression driven by PM split-GAL4 drivers in the five major intestinal regions (R1–R5) stained for GFP (anti-GFP in green), progenitor cells (anti-HRP in red), enteroendocrine cells (anti-Pros in white), and nuclei (DAPI in blue). (G) Graph showing percent of progenitor cells, EEs, ECs, and all cells labelled by *PM-AD* and *PM-DBD* driving *UAS-GFP.nls* expression in each of the five intestinal regions. Progenitor cells were defined as cells showing anti-HRP staining, EEs as cells showing anti-Pros staining, and ECs as cells lacking both anti-HRP and anti-Pros staining. All cells showed DAPI staining. Cells were counted from representative fields of view for each region from 5–6 intestines. Graph shows means with SD. Scale bars; 500μm (A–E), 25μm (F). Complete genotypes are listed in Table S2.

## MATERIALS AND METHODS

### Drosophila strains and husbandry

All fly strains were cultured on standard Bloomington media (https://bdsc.indiana.edu/information/recipes/bloomfood.html) at 25°. The full genotypes of all starting stocks used in this study are listed in Table S1, and the drivers and responders used in each figure are listed in Table S2. The *PBac{UAS-DSCP-6XEGFP}VK00018* and *PBac{UAS-DSCP-6XEGFP}attP2* strains were gifts from Steve Stowers (Montana State University).

### Transgenes

Transgene construction was performed using NEBuilder HiFI DNA Assembly Master Mix (New England Biolabs, Catalog E2621), restriction enzyme-digested plasmids, and PCR products generated with primers listed in Table S3. Plasmids were sequence verified, amplified, and sent to Rainbow Transgenic Flies (Camarillo, CA) for injection.

### Pan-midgut split-GAL4 transgenes

The *Zip-GAL4DBD* and *p65AD-Zip* open reading frames were assembled downstream of the 1kb *CG10116* enhancer fragment by combining the XhoI/XbaI-digested *CG10116p-KD::PEST* plasmid (Buddika et al., 2020b) with either a *Zip-GAL4DBD*-containing PCR fragment amplified with oligo pair 3855/3856 (see Table S3 for oligo sequences) from *pBPZpGAL4DBDUw* or a *p65AD-Zip*-containing PCR fragment amplified with oligo pair 3857/3858 from *pBPp65ADZpUw* (both plasmids (Addgene 26233 and 26234) were gifts from Gerald Rubin (Pfeiffer et al., 2010)). Transgenesis of these plasmids yielded *P{CG10116-GAL4*.*DBD}su(Hw)attP6* and *P{CG10116-p65*.*AD}attP40*.

### Cell type split-GAL4 reference lines

Cell type-specific split-GAL4 reference lines were generated by swapping the *CG10116* enhancer fragment in the plasmids described above for either the 2.4kb *GMR57F07* enhancer fragment from *Dh31* or the synthetic *GBE* fragment containing tandem arrays of Grainyhead and Suppressor of Hairless binding sites (Beehler-Evans and Micchelli, 2015; Furriols and Bray, 2001; Guo et al., 2013). The *GMR57F07* enhancer fragment was amplified from genomic DNA using oligo pair 3955/3956, while the *GBE* sequence was amplified from the previously described *3Xgbe-smGFP::V5::nls* plasmid (Buddika et al., 2020a) with oligo pair 3450/3451. Transgenesis of these plasmids yielded *P{R57F07-p65*.*AD*.*A}attP40, P{R57F07-GAL4*.*DBD*.*A}attP2*, and *P{GBE-GAL4*.*DBD}attP2*.

### *Enhancer-GAL4* and *-smGFP.V5.nls* transgenes

Enhancers were PCR amplified using oligonucleotide primers listed in Table S3. These enhancers were cloned via Gibson Assembly into *3Xgbe*-excised versions of either the *3Xgbe-smGFP::V5::nls* plasmid (Buddika et al., 2020a) or the same plasmid in which *smGFP::V5::nls* had been replaced with *GAL4*.

### Dissections and immunostaining

For most analyses of *UAS-6XGFP* and *UAS-Stinger* expression, gastrointestinal tracts were dissected in 1⨯ PBS (137mM NaCl, 2.7mM KCl, 10mM Na_2_HPO_4_, KH_2_PO_4_, pH 7.4), fixed in 4% paraformaldehyde (Electron Microscopy Sciences, Cat. No. 15714) in PBS for 45 min, washed multiple times in 1xPBT (1xPBS, 0.1% Triton X-100), including one wash with 5μg/ml DAPI in PBT, and mounted in DABCO-containing mounting medium. For all antibody labeling except Delta staining, samples were fixed as above, and then incubated at 4° overnight with primary antibodies, including mouse anti-Prospero (MR1A, Developmental Studies Hybridoma Bank, 1:100), mouse anti-V5 (MCA1360GA, Bio-Rad, 1:250), and rabbit anti-GFP (A11122, Life Technologies, 1:1000). The following day, samples were washed in 1xPBT and incubated for 2–3 hours with secondary antibodies, including AlexaFluor 488- and 568-conjugated goat anti-rabbit, -mouse, -rat and -chicken antibodies (Life Technologies, 1:1000). AlexaFluor 647-conjugated goat-HRP antibodies were used in the secondary antibody solution whenever required. Finally, samples were washed multiple times in 1xPBT, including one wash with 5μg/ml DAPI in PBT, and mounted in Vectashield mounting medium (Vector Laboratories). An alternative staining protocol was used for mouse anti-Delta (C594.9B, Developmental Studies Hybridoma Bank, 1:500) staining as described in (Buddika et al., 2020b) and these samples were mounted in ProLong Diamond mounting medium (Invitrogen, P36970). Intestines stained with anti-Delta antibody were also stained with anti-GFP antibody, since the methanol steps required in this protocol quenched GFP fluorescence.

### Microscopy and image processing

Images of whole flies and dissected intestines during the screens were collected on a Zeiss Axio Zoom microscope. Images of immunostained intestines were collected on a Leica SP8 Scanning Confocal microscope. Samples to be compared were collected under identical settings on the same day, image files were adjusted simultaneously using Adobe Photoshop, and figures were assembled using Adobe Illustrator.

### Expression pattern analysis

Intestines from 4- to 8-day old flies were analyzed and the expression patterns of at least five independent intestines were scored per genotype. For the primary screen, third instar larvae and adults were examined for fluorescence. When fluorescence was observed in adults, males and females were dissected and expression patterns were characterized by scoring the presence or absence of GFP expression in each of the five main intestinal regions (R1–R5). Variability among samples was noted. For the secondary screen, males and females were examined for drivers where a dimorphism was noted in the initial screen; only females were examined for the remainder.

Expression patterns were characterized by semiquantitatively estimating the number of cells that displayed GFP expression in each of 11 intestinal subregions (R1A, R1B, R2A, R2B, R2C, R3, R4A, R4B, R4C, R5A, and R5B) (Buchon et al., 2013) on a scale from 0 to 3, where 0 indicated no cells, 1 indicated some cells (∼1–49%), 2 indicated most cells (∼50–99%), and 3 indicated all cells (∼100%). The variability in GFP intensity among cells of labeled intestines was also analyzed semiquantitatively: given a 0 when no midgut expression was present, a 1 when there was more than twofold difference among at least 25% of cells, and a 2 when all labeled cells were roughly the same intensity. The variability of expression among intestines of the same genotype was scored on a scale from 0 to 3, where 0 indicated that all intestines had the same pattern, 1 indicated that the majority had the same pattern, 2 indicated that there was a pattern common to all samples but also some notable differences among them, and 3 indicated that all intestines displayed different patterns.

### Data Availability

Plasmids are available upon request. Extant stocks are available from the Bloomington Drosophila Stock Center or upon request. Fig S1 and S2 show the results of control experiments. Fig S3 shows representative images summarized in Table 2. Table S1 is the reagent table. Table S2 lists the genotypes of all strains shown in Figures. Table S3 is a list of all oligonucleotide primers. Table S4 lists expression patterns of new *GAL4* and *GFP* transgenes. File S1 describes driver characterizations in the primary screen. File S2 lists expression patterns characterized in the secondary screen. The results of our primary and secondary screens are also available online at https://bdsc.indiana.edu/stocks/gal4/midgut_splitgal4.html. File S3 lists drivers with clear regional expression boundaries. The authors affirm that all data necessary for confirming the conclusions of the article are present within the article, figures, and tables.

**Table 1.** Driver characterization in primary screen

|  | **GAL4DBD** | **p65AD** |
| --- | --- | --- |
| **Drivers screened** | 4,296 | 3,008 |
| **Intestinal expression** | 417 | 173 |
| **Adult** | 302 | 122 |
| **Sex-dependent** | 21 | 16 |
| **Stage-dependent** | 257 | 101 |
| **Larva only** | 115 | 51 |
| **Adult only** | 142 | 50 |
| **Regional** | 100 | 42 |
| **Variable expression** | 47 | 23 |

**Table 2.** *UAS-GFP* expression in anterior, middle and posterior regions when reference drivers were combined^*a*^

|  | *PM-DBD* | *P-DBD* | *EE-DBD* | *EC-DBD* |
| --- | --- | --- | --- | --- |
| *PM-AD* | + – + | + + + | – – + | + – + |
| *P-AD* | + – + | + – + | – – – | – – – |
| *EE-AD* | – – + | – – – | + + + | – – – |
| *EC-AD* | + – + | – – – | – – – | Lethal |

<sup>a</sup> Presence (+) or absence (–) of GFP expression indicated for anterior (R1–R2), middle (R3) and posterior (R4–R5) regions.

## RESULTS

### Construction and verification of reference drivers for primary screen

To screen the split-GAL4 collection efficiently, we first designed reference stocks to restrict expression of individual split drivers to the midgut. These reference stocks contained an intestine-specific split-GAL4 transgene (encoding either the p65 activation domain (p65AD) or GAL4 DNA-binding domain (GAL4DBD)) and a clearly detectable *UAS-reporter* transgene. We cloned the *p65AD* or *GAL4DBD* open-reading frame downstream of an enhancer from the *CG10116* gene, whose activity in other transgenic contexts was limited to the gastrointestinal tract (Buddika et al., 2020b). To verify that these new *CG10116-p65AD* and *CG10116-GAL4DBD* drivers were expressed throughout the intestine as expected, we examined flies carrying both transgenes and either *P{20XUAS-6XGFP}attP2* or *P{20XUAS-DCSP-6XEGFP}attP2*, which express a hexameric version of green fluorescent protein under the control of 20 copies of *UAS* (Shearin et al., 2014; Williams et al., 2019). Because these insertions differ only in promoter sequences and were interchangeable in marking intestinal cells, we will refer to them both as *UAS-6XGFP*. Both larvae and adults displayed clear, strong intestinal fluorescence, which could be seen through the abdominal cuticle, and little fluorescence elsewhere (Fig 1B, C). All midgut regions from both females and males were strongly fluorescent except for region 3 (R3), which was much dimmer (Fig 1D, E). Importantly, we did not see *UAS-6XGFP* expression when *CG10116-p65AD* or *CG10116p-GAL4DBD* were combined respectively with “empty” GAL4DBD or p65AD transgenes carrying no enhancer sequences (Fig S1). These results indicated that we could use these new drivers, which we refer to as *PM-AD* (*pan-midgut-p65AD*) and *PM-DBD* (*pan-midgut*-*GAL4DBD*) in the remainder of the text, to identify split-GAL4 drivers that have midgut expression, with the possible exception of drivers that express only in R3.

To evaluate the intestinal cell types expressing *PM-AD* and *PM-DBD* in detail, we combined these drivers with a nuclear GFP reporter (*UAS-GFP.nls*) that was easier to score on a cell-by-cell basis than *UAS-6XGFP*. We detected nuclearly localized GFP expression with an anti-GFP antibody and counterstained intestines with anti-horseradish peroxidase (HRP) and anti-Prospero (Pros) antibodies and DAPI, which label progenitor cells, EEs, and all cells, respectively (Micchelli and Perrimon, 2006; O’Brien et al., 2011; Shiga et al., 1996). Progenitor cells include both ISCs and EBs, two cell types that have highly similar transcriptional profiles (Dutta et al., 2015; Hung et al., 2020). While not directly labeled, we classified ECs as cells with neither anti-HRP nor anti-Pros antibody staining. We found that most cells in the intestinal epithelium expressed GFP, ranging from 75% in R3 to 94% in R2 and R4 (Fig 1F, G). This included almost all ECs (85–100%) and the majority of progenitor cells (65–97%). EEs in most regions, however, were not efficiently marked. While 68% were labeled in R2, only 5% were labeled in R5 and none were labeled in R3. These results indicated that the *CG10116* enhancer was active in the majority of intestinal cells and that, consequently, *PM-AD* or *PM-DBD* could be paired with any other split-GAL4 driver to demonstrate intestinal expression with the possible exception of drivers expressed in only a subset of EEs.

We noted that GFP expression was affected by the choice of *UAS* construct. *P{20XUAS-6XGFP}* showed high non-intestinal expression with *PM-DBD*, while *P{20XUAS-DSCP-6XEGFP}* did not. Within the intestine, we saw no differences in the patterns of GFP fluorescence with either *UAS-6XGFP* construct, *UAS-GFP.nls* or *UAS-Stinger*, which expresses a nuclear GFP variant (Barolo et al., 2000) (Fig S2), although each construct showed a characteristic expression level (with *GFP.nls* expression low enough to require anti-GFP antibody staining for reliable detection). There was a single idiosyncratic exception: only *UAS-GFP.nls* was strongly detected in R3 in the presence of PM drivers. The general interchangeability of *UAS-GFP* responders in the intestine meant that we could choose the one with optimal expression for each experimental approach.

### Primary screen identifies 424 split-GAL4 drivers with adult midgut expression

We used *PM-AD* or *PM-DBD* in combination with *UAS-6XGFP* to screen the Bloomington Drosophila Stock Center collection of split-GAL4 stocks, which carry driver transgenes generated at Janelia Research Campus and the Institute of Molecular Pathology in Vienna (Dionne et al., 2018; Tirian and Dickson, 2017). From the 7,304 stocks screened, we identified 590 total drivers, including 417 GAL4DBD and 173 p65AD drivers, giving abdominal fluorescence in third instar larvae, young female adults, and/or young male adults (Table 1). A total of 424 of these included adult midgut expression. As a control for our detection method, we randomly chose 50 drivers that were scored as negative in whole animals and verified that dissected intestines lacked GFP fluorescence. However, we cannot rule out the possibility that our approach failed to detect drivers that sparsely or weakly labeled intestines. Among the 590 positive drivers, we noted several expression categories, including drivers that labeled subsets of cells based on sex, cell morphology, stage, and/or region (Fig 2, Table 1, File S1). The 37 sex-dependent drivers included six expressed only in adult males, seven expressed only in adult females, and 24 expressed in distinct patterns in each sex (Fig 2A). Cell morphology-specific drivers labeled subsets of cells of similar appearance throughout the adult midgut (Fig 2B). (Cell types were determined systematically in the subsequent secondary screens.) Stage-specific drivers were expressed in either larvae or adults (Fig 2C). Finally, region-dependent drivers selectively labeled patches of cells along the anterior–posterior (A–P) axis of the adult intestine, and ranged in expression from a small number of cells in a subregion to half of the gut (Fig 2D). An additional 70 drivers displayed variable expression patterns among adult intestines. We noted this variability (File S1) and included these drivers in subsequent analyses.

**Figure 2.**
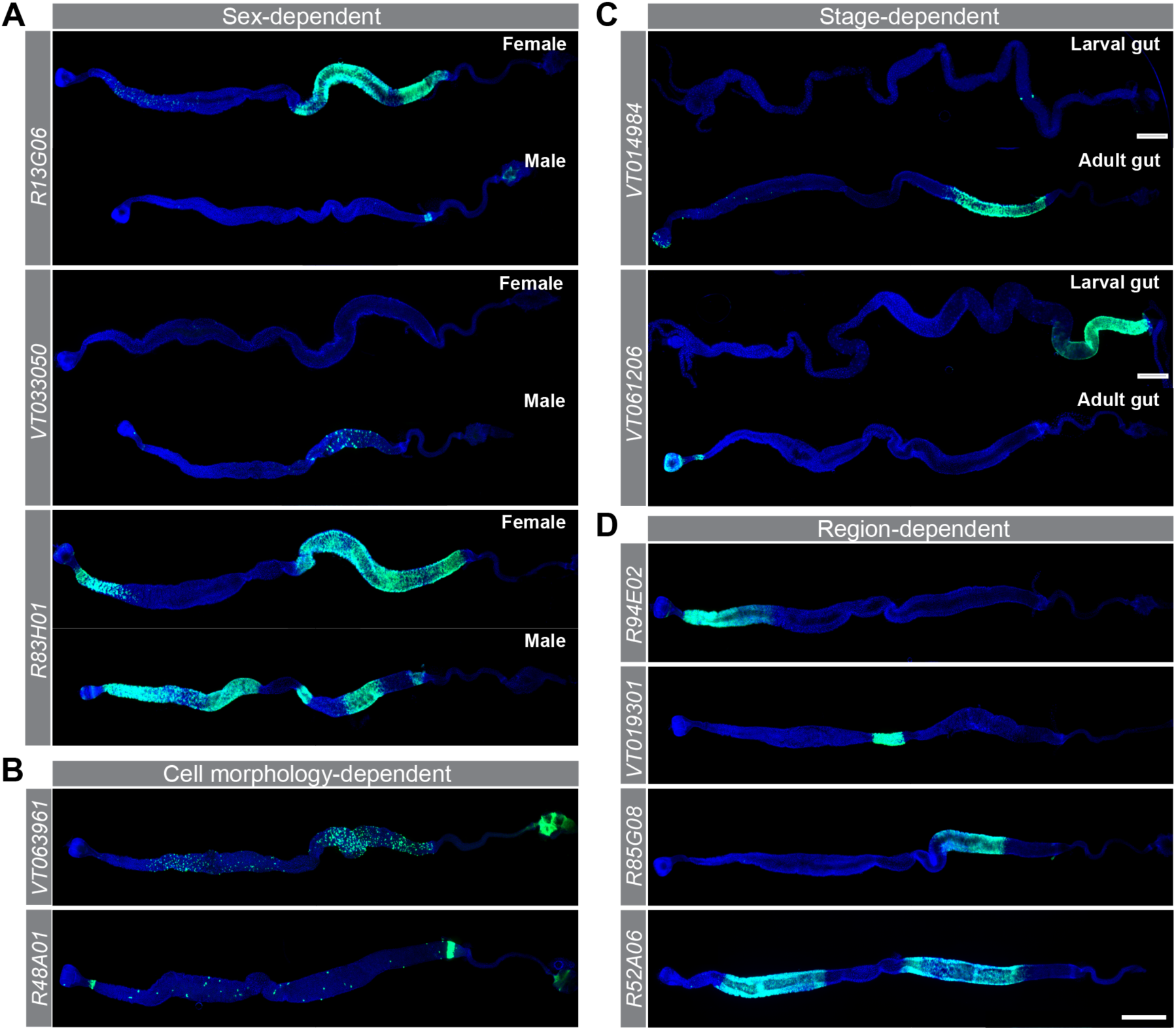
The primary screen identified a variety of split-GAL4 expression patterns in the intestine. These micrographs show intestinal *UAS-6XGFP* expression (green) when split-GAL4 drivers were combined with PM drivers and stained with DAPI (blue). (A) Examples of sex-dependent patterns. *R13G06-p65AD* drives expression only in females and *VT033050-GAL4DBD* only in males. *R83H01-GAL4DBD* drives expression in different patterns in females and males. (B) Examples of cell morphology-specific patterns. *VT063961-GAL4DBD* and *R48A01-GAL4DBD* drive expression in similar-looking cells in distinctly different patterns. (C) Examples of stage-dependent midgut patterns. *VT014984-GAL4DBD* drives expression only in adults and *VT061206-GAL4DBD* only in larvae (D) Examples of region-dependent patterns including anterior (*R94E02-p65AD*), middle (*VT019301-GAL4DBD*), posterior (*R85G08-GAL4DBD*), or both anterior and posterior (*R52A06-GAL4DBD*) expression. Scale bars; 500μm. Complete genotypes are listed in Table S2.

### Split-GAL4 reference strains label distinct intestinal cell types

To identify the cell types labeled by the 424 split-GAL4 drivers with adult expression, we assembled a collection of p65AD and GAL4DBD reference drivers for each of the three main midgut cell types: progenitor cells, EEs, and ECs. For the progenitor cells and ECs, we used drivers identified in the primary screen (*VT004241-p65AD* and *VT024642-GAL4DBD* for progenitor cells; *VT004958-p65AD* and *VT004958-GAL4DBD* for ECs). For EEs, we generated new drivers containing the *GMR57F07* enhancer fragment, which was previously shown to drive GAL4 expression in EEs (Beehler-Evans and Micchelli, 2015). Because the extent and specificity of expression of these drivers was critical to their utility, we rigorously benchmarked their cell type expression using the *UAS-GFP* reporters and anti-HRP and anti-Pros staining described above. *VT004241-p65AD* and *VT024642-GAL4DBD* combined with *UAS-Stinger* labeled 97–100% of progenitor cells in R1, R2, R4, and R5 and 63% in R3 (Fig 3A, E). Driver expression was specific to progenitor cells, since only 4% of GFP-positive cells lacked anti-HRP staining. *R57F07-p65AD* and *R57F07-GAL4DBD* combined with *UAS-6XGFP* labeled 52–57% of EEs in R1, R2, R3 and R5, but only 3% in R4 (Fig 3B, F). Although these drivers did not label all EEs, we adopted them as reference drivers since they labeled more EEs than any other driver tested (including a split-GAL4 version of the widely used *P{GawB}pros*^*V1*^ driver, which labels most EEs, that we generated using the HACK method (Balakireva et al., 1998; Lin and Potter, 2016)). Because the EC drivers, *VT004958-p65AD* and *VT004958-GAL4DBD*, caused lethality when combined, we evaluated their expression using the *PM-AD* and *PM-DBD* drivers with *UAS-GFP.nls*. Both *EC* drivers labeled 98–100% of ECs in all regions except in R3, where *VT004958-p65AD* labeled 93% and *VT004958-GAL4DBD* labeled 77% of ECs, with almost no other cells labeled (Fig 3C, D, G, H). Collectively, this characterization indicated that these six drivers showed the expected cell type expression. We refer to these drivers by their cell (*P-, EE-*, or *EC-*) and split type (*-AD* or *-DBD*) below, where we use them in combination with *UAS-6XGFP*.

**Figure 3.**
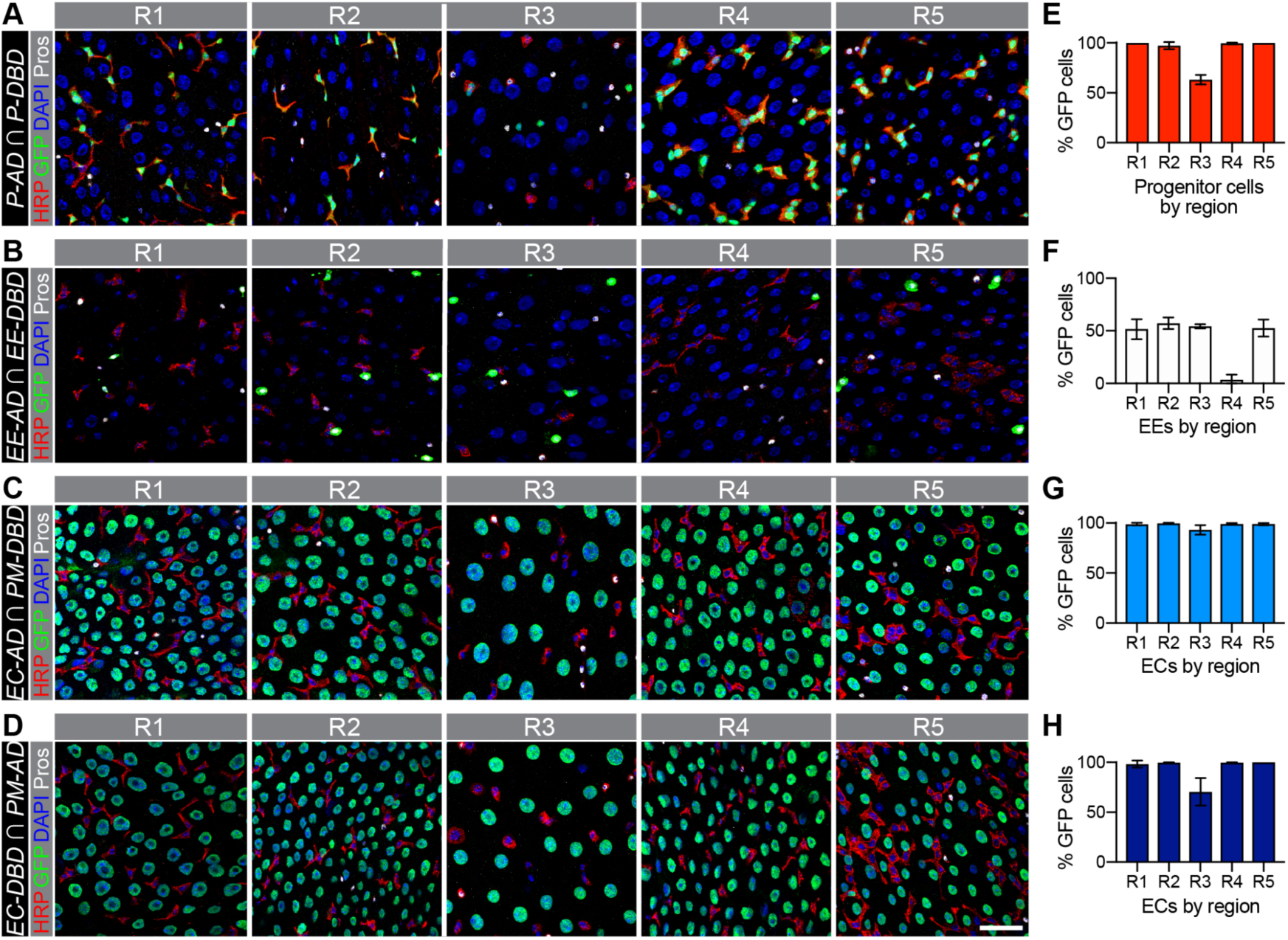
Reference split-GAL4 drivers label specific intestinal cell types. (A–D) GFP expression in intestinal regions R1–5 driven by (A) progenitor (P) drivers (*VT004241-p65AD* ∩ *VT024642-GAL4DBD*), (B) EE drivers (*R57F07-p65AD* ∩ *R57F07-GAL4DBD*), (C) an *EC-AD* driver (*VT004958-p65AD* ∩ *CG10116-GAL4DBD*), and (D) an *EC-DBD* driver (*VT004958-GAL4DBD* ∩ *CG10116-p65AD*). Anti-GFP antibody staining is shown in green with counterstaining for progenitor cells (anti-HRP in red), EEs (anti-Pros in white), and all cell nuclei (DAPI in blue). Scale bar; 25 μm. (E–H) Histograms showing the percent of (E) progenitor cells, (F) EEs, and (G,H) ECs labeled with GFP by region. Cell identities were defined and cells counted as in Figure 1. Complete genotypes are listed in Table S2.

To further validate the cell specificity of these reference drivers, we made pairwise combinations of the reference drivers (Table 2, Fig S3). For the most part, the results were as expected: no GFP was detected when reference drivers expressed in distinct cell types were combined with each other, but GFP was detected when reference drivers labeling the same cell types were combined. We noted two exceptions. First, GFP was poorly expressed in R3 with the PM drivers, as noted above, as well as with the P and EC drivers. Second, we detected GFP expression in some large cells in R5 and occasionally R4 with the P and EE drivers, suggesting driver expression in a few ECs, which are the largest intestinal cells. Nevertheless, these results confirmed the specificity and effectiveness of the reference drivers in most regions.

### Secondary screen defines cell type expression of 424 split-GAL4 drivers

To define the cell type expression patterns of the 424 split-GAL4 transgenes identified in the primary screen that drive in adults, we combined each with P, EE, and EC reference drivers and *UAS-6XGFP*. Expression patterns were analyzed in females for all drivers, and in males for drivers with sex-dependent expression in the primary screen. We semiquantitatively scored the intensity and variability of expression in the 11 subregions along the midgut A–P axis (R1A, R1B, R2A, R2B, R2C, R3, R4A, R4B, R4C, R5A, and R5B) (File S2). We identified 177 drivers with progenitor cell expression, 116 with EE expression, and 332 with EC expression (Table 3). While many of the EC-expressing drivers expressed only in ECs (155), fewer lines displayed progenitor cell- or EE-specific expression (36 and 4, respectively). Many cell type-specific drivers labeled cells in only a specific subregion or in a subset of cells throughout the midgut (Fig 4A– D). In contrast, 122 lines displayed expression in some combination of two cell types, and 63 lines displayed expression in all three cell types. By comparing driver expression profiles in the presence of all four reference drivers, we could deconstruct some complex patterns seen in our primary screen into combinations of simpler cell type-specific patterns (Fig 4E–H). We also identified drivers that labeled two or three cell types, each displaying a distinct regional pattern (Fig 4E–G). A total of 81 drivers (19%) displayed some variability in expression among the intestines of a single sample, but, interestingly, no driver showed variable expression with all reference drivers, suggesting that the expression variability was cell type dependent.

**Table 3.**
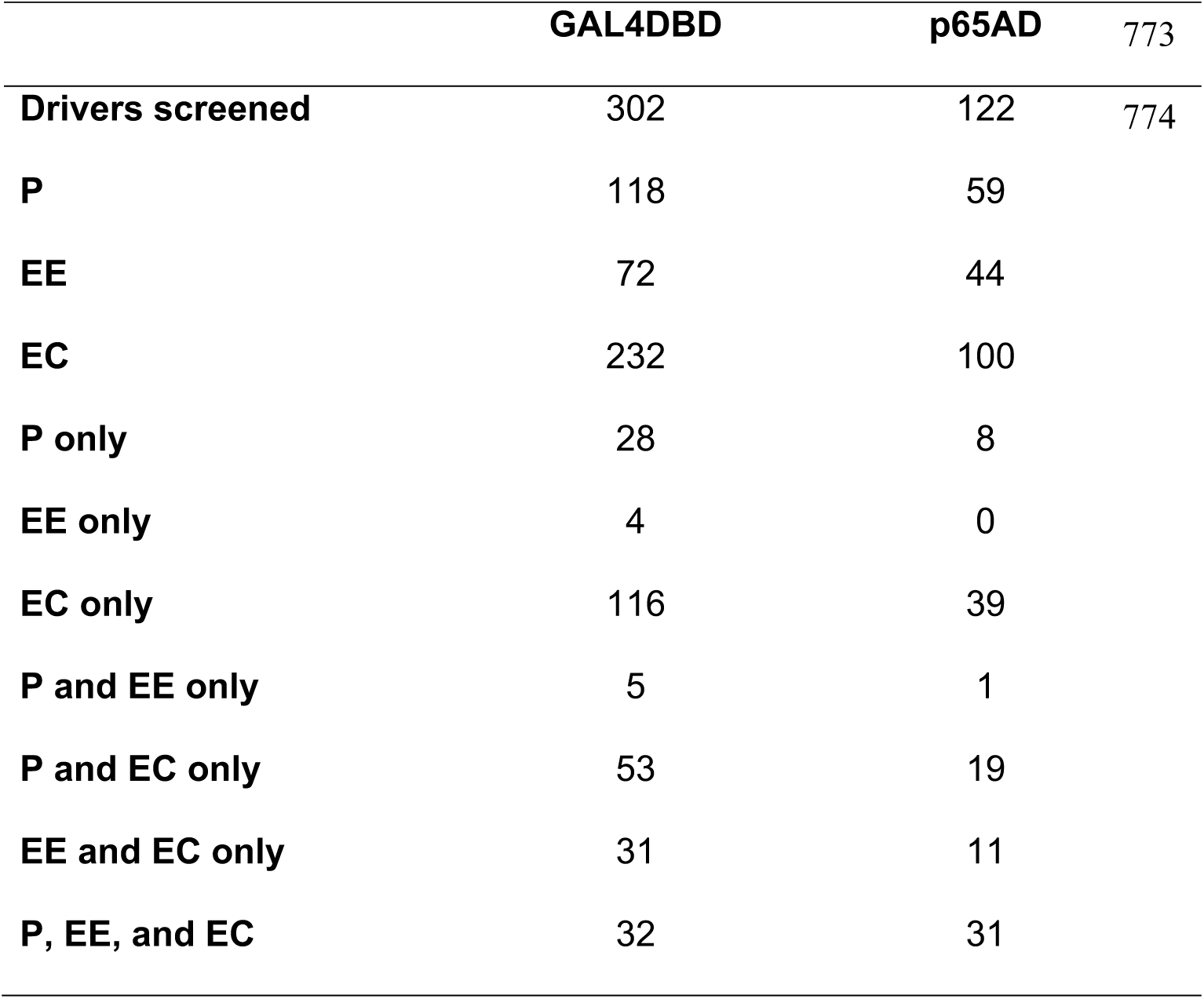
Cell type expression of drivers in secondary screen

**Figure 4.**
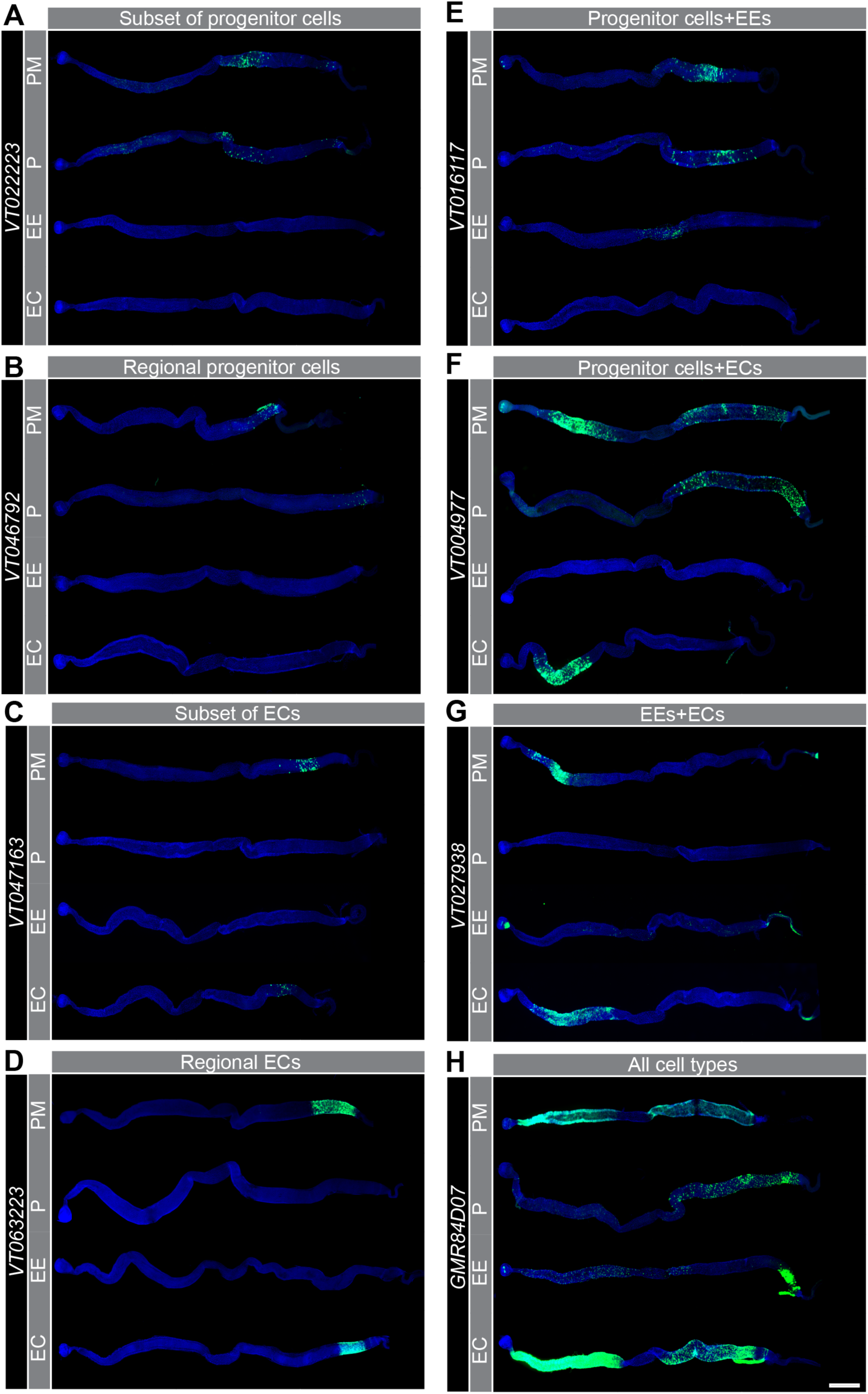
Cell type expression of representative split-GAL4 drivers. (A–H) Intestinal expression of *UAS-6XGFP* (green) when DBD drivers are combined with *PM-, P-, EE-*, and *EC-AD* drivers. (A) *VT022223-GAL4DBD* drives expression in a subset of progenitor cells throughout the midgut, (B) *VT046792-GAL4DBD* in a subset of R5 progenitor cells, (C) *VT047163-GAL4DBD* in a subset of R4 ECs, (D) *VT063223-p65AD* in most R5 ECs, (E) *VT016117-GAL4DBD* in progenitor cells and EEs, (F) *VT004977-GAL4DBD* in anterior ECs and posterior progenitor cells, (G) *VT027938-GAL4DBD* in ECs and EEs, and (H) *R84D07-GAL4DBD* in progenitor cells, ECs, and EEs. Intestines were counterstained with DAPI (blue). Scale bar; 500 μm. Complete genotypes are listed in Table S2.

We also used the cell type-specific data to further examine the drivers with sex-dependent expression profiles identified with the PM reference drivers (File S2). Of the 24 drivers showing distinctly different expression patterns in males and females, differences in 12 of these drivers could be explained by EC expression alone, while dimorphisms in another three drivers could be explained by expression in ECs in combination with other cell types (EE, P, or both). With three other drivers, we saw no expression with the P, EE, or EC reference drivers in either sex, and, with three drivers, we saw no cell type clearly responsible for the sex-dependent expression observed in our primary screen. Altogether, this analysis provides a comprehensive resource of split-GAL4-dependent expression profiles that can be used to target cells in a region- and cell type-specific manner. A database of expression profiles can be viewed at https://bdsc.indiana.edu/stocks/gal4/midgut_splitgal4.html.

### Split-GAL4 drivers label subsets of progenitor cells

To extend classification of the split-GAL4 drivers, we prepared an additional split-GAL4 reference driver that expresses GAL4DBD under the control of the synthetic *GBE* enhancer, known to direct expression in EBs, but not ISCs (Furriols and Bray, 2001). To verify expression of this *EB-DBD* driver, we combined it with *PM-AD* and *UAS-GFP.nls* and stained intestines with fluor-tagged antibodies against GFP, HRP, and Pros as well as Delta (Dl) protein, which is an established marker of ISCs (Fig 5A) (Micchelli and Perrimon, 2006). We classified EBs as cells detected by anti-HRP antibodies, but not anti-Dl antibodies. We found that 82–99% of EBs expressed GFP in R1, R2, R4, and R5 (very few R3 cells were labeled), but only 6–13% of non-EB cells expressed GFP, indicating that *EB-DBD* effectively labels the EB subpopulation of progenitor cells in most regions (Fig 5B). We then screened the 59 p65AD drivers that had shown progenitor cell expression and found 44 with EB expression (File S2). Some of these drivers labeled EBs predominantly in one region, such as *P{R78B06-p65.AD}* and *P{VT055900-p65.AD}*, while some labeled EBs throughout the intestine, such as *P{R10F10-p65.AD}* (Fig 5C–E). We also fortuitously identified two drivers with EB expression that were not identified as expressing in progenitor cells with *P-AD* or *P-DBD*. To complement this EB analysis, we attempted to generate *EB-AD* and ISC reference drivers, but they were inviable or not specific to the targeted cell type.

**Figure 5.**
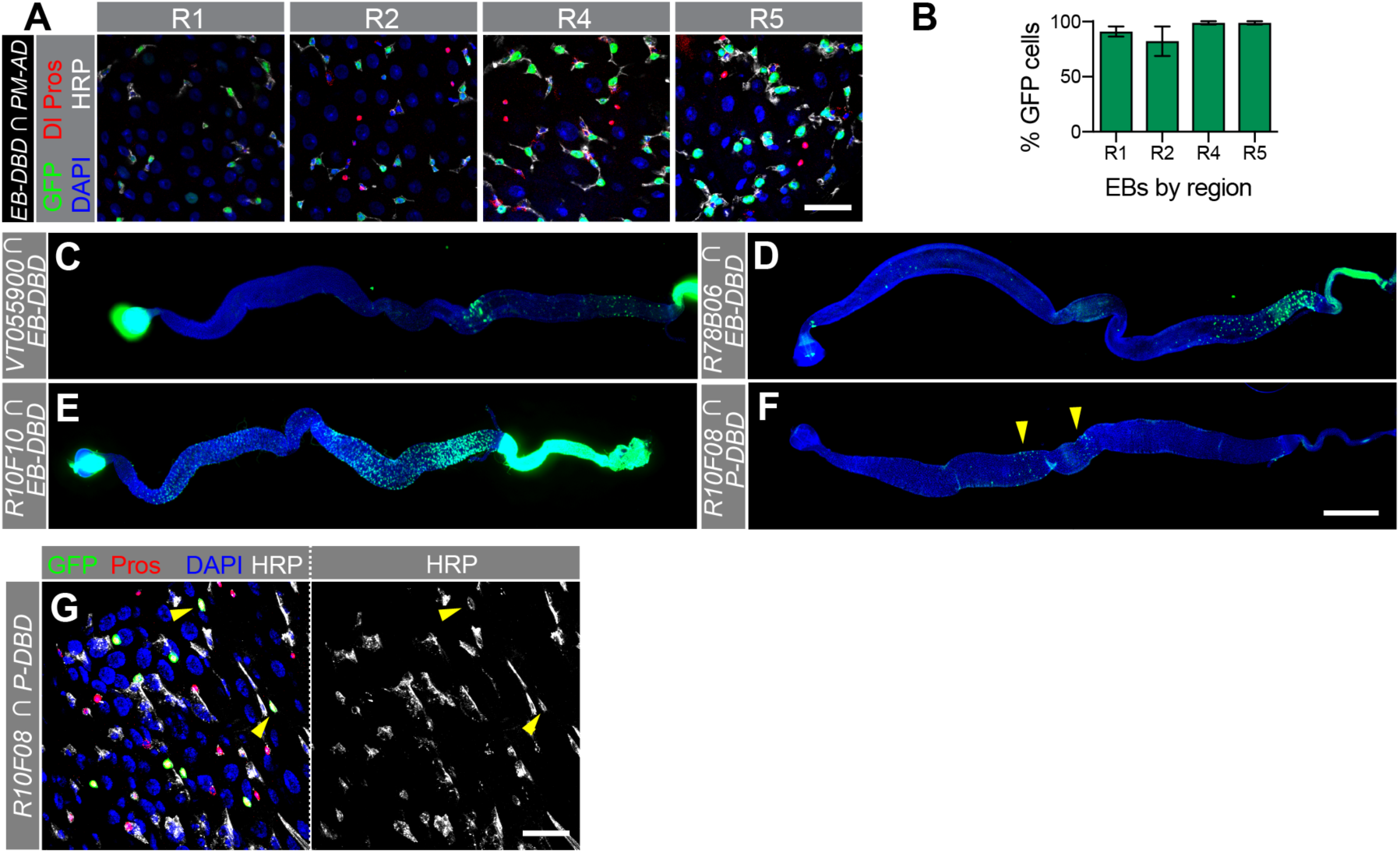
The secondary screen identified split-GAL4 drivers that label specific progenitor and EE cell populations. (A) GFP expression in R1, R2, R4, and R5 driven by *EB-DBD* combined with *PM-AD*. Anti-GFP antibody staining is shown in green with counterstaining for ISCs (anti-Dl, red cytoplasmic staining), EEs (anti-Pros, red nuclear staining), progenitors (anti-HRP, white), and the DNA marker DAPI (blue). (B) Quantification of the percent of EBs labeled by GFP in four intestinal regions. EBs are defined as cells showing anti-HRP staining but not anti-Dl staining. (C, D) Intestinal expression of *UAS-6XGFP* (green) when EB-DBD is combined with *VT055990-p65AD* or *R78B06-p65AD*. (E, F) Intestinal expression of *UAS-Stinger* when *R10F08-p65AD* is combined with *PM-DBD* (E) or *P-DBD* (F, yellow arrowheads indicate locations of small subsets of labeled cells). (G) *UAS-Stinger* expression driven by *R10F08-p65AD* and *PM-DBD* in an intestine counterstained for EEs (anti-Pros, red), progenitors (anti-HRP, white), and DAPI (blue). Yellow arrowheads indicate HRP+, Pros+, Stinger+ cells. Scale bars: 25μm (A, G), 500 μm (C-F). Complete genotypes are listed in Table S2.

### Identification of a split-GAL4 driver that specifically labels a novel subtype of EEs

In an attempt to identify subpopulations of ISC cells, we focused on the 15 p65AD drivers that showed GFP expression with *P-DBD*, but not *EB-DBD*. We observed that two drivers labeled a small population of small cells in the central midgut. One driver, *R10F08*-*p65AD*, restricted GFP expression to these cells when combined with *P-DBD* (Fig 5F). To determine the cell type identity of this cell population, we combined *R10F08*-*p65AD* with *P-DBD* and *UAS-Stinger* and stained intestines with antibodies against HRP and Pros. Surprisingly, we found that the majority of cells that expressed *UAS-Stinger* were labeled by both anti-HRP, a progenitor cell marker, and anti-Pros, an EE marker (Fig 5G). Although these two markers are mutually exclusive in most intestinal cells, Hung et al. (2020) recently identified a subclass of EEs with similar location and morphology that also express EE and progenitor cell markers. We conclude that *R10F08*-*p65AD* in combination with *P-DBD* labels this novel EE subtype, demonstrating that split-GAL4 drivers can be used to identify novel cell populations with distinct molecular properties.

### Split-GAL4 combinations label regional cell types and subregions

As described above, the expression of many drivers is limited to specific intestinal regions. We reexamined these regional drivers and identified 48 p65AD and 62 GAL4DBD drivers that have well-defined expression boundaries along the A–P axis (File S3). Figure 6A shows five EC drivers expressing in progressively more posterior locations within R1 and R2. It was clear from such overlapping patterns that we should be able to identify driver pairs defining smaller cell populations. Figure 6B shows that two EC drivers expressed in the posterior midgut (*VT004417-p65AD* and *VT045598-GAL4DBD*) could be combined to define a narrower band of cells with a particularly distinct anterior boundary in R4. Figures 6C and D show other examples. Additional studies are needed to determine the functional significance of discrete cell populations defined by driver pairs, but our database of driver expression patterns (https://bdsc.indiana.edu/stocks/gal4/midgut_splitgal4.html) provides information for predicting which driver combinations might be experimentally useful for delineating new intersectional patterns.

**Figure 6.**
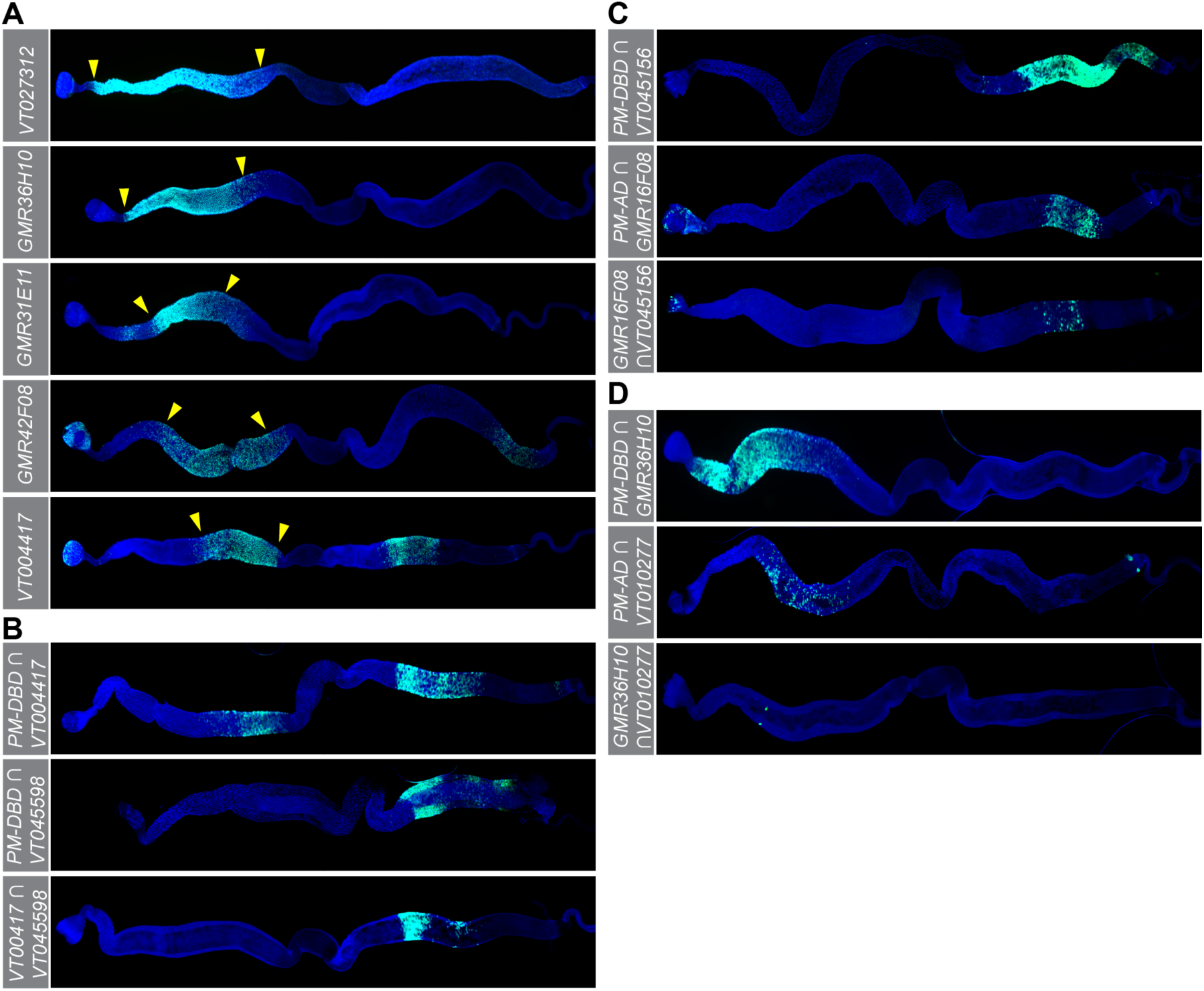
The expression of split-GAL4 drivers can display clear regional boundaries and, when combined, drivers can label small subsets of cells. (A) Intestinal expression of *p65AD* drivers in anterior regions demonstrated in combination with *PM-DBD*. Other tests showed these five drivers are expressed solely in ECs. Yellow arrowheads indicate regional boundaries. (B) *VT004417-p65AD* and *VT045598-GAL4DBD* are expressed broadly in the posterior intestine as shown in combinations with PM drivers, but they define a more restricted expression pattern when combined with each other. (C–D) Similarly, other driver pairs define discrete cell populations. *UAS-6XGFP* expression shown in green; DAPI staining in blue. Complete genotypes are listed in Table S2.

### Identification of midgut enhancers

To address the lack of molecularly defined enhancers with adult midgut activity, we tested whether enhancer fragments from the split-GAL4 transgenes identified above could drive similar expression in other transgenic contexts. We cloned ten enhancer fragments into a transformation vector encoding full-length GAL4 and saw seven examples where the new GAL4 driver induced *UAS-Stinger* expression in a similar, although not always identical, pattern as a split-GAL4 driver with the same fragment combined with *PM-AD* or *PM-DBD* (Fig 7A, Table S4). Figure 7A shows that the *VT043613* and *GMR61H08* fragments drove almost identical expression in both systems, while *GMR16G08, GMR86G08*, and *GMR28G08* drove expression in both systems with relatively minor regional differences. In addition, *VT024642* drove expression in progenitor cells throughout the midgut in both systems (shown in Fig 7A and quantified for the *GAL4* version in Fig 7B, C). While these examples showed enhancer fragments can drive similar patterns in different transgenic contexts, we saw three enhancer fragments that failed to replicate the split-GAL4 pattern in the *GAL4* vector and six out of eleven fragments that failed when transferred to a *smGFP.V5.nls* vector (Table S4). Others have seen that expression patterns are not always recapitulated when enhancer fragments are transferred among vectors (Chen et al., 2019; Dionne et al., 2018). Such discrepancies may reflect regulatory interactions between transgene sequences or between transgene and adjacent genomic sequences. Regardless of the reasons for inconsistency, our analysis shows that enhancer sequences characterized using the split-GAL4 system can be used successfully to generate new transgenic tools for labeling and manipulating defined intestinal cells and highlights the value of our extensive documentation of split-GAL4 driver expression patterns.

**Figure 7.**
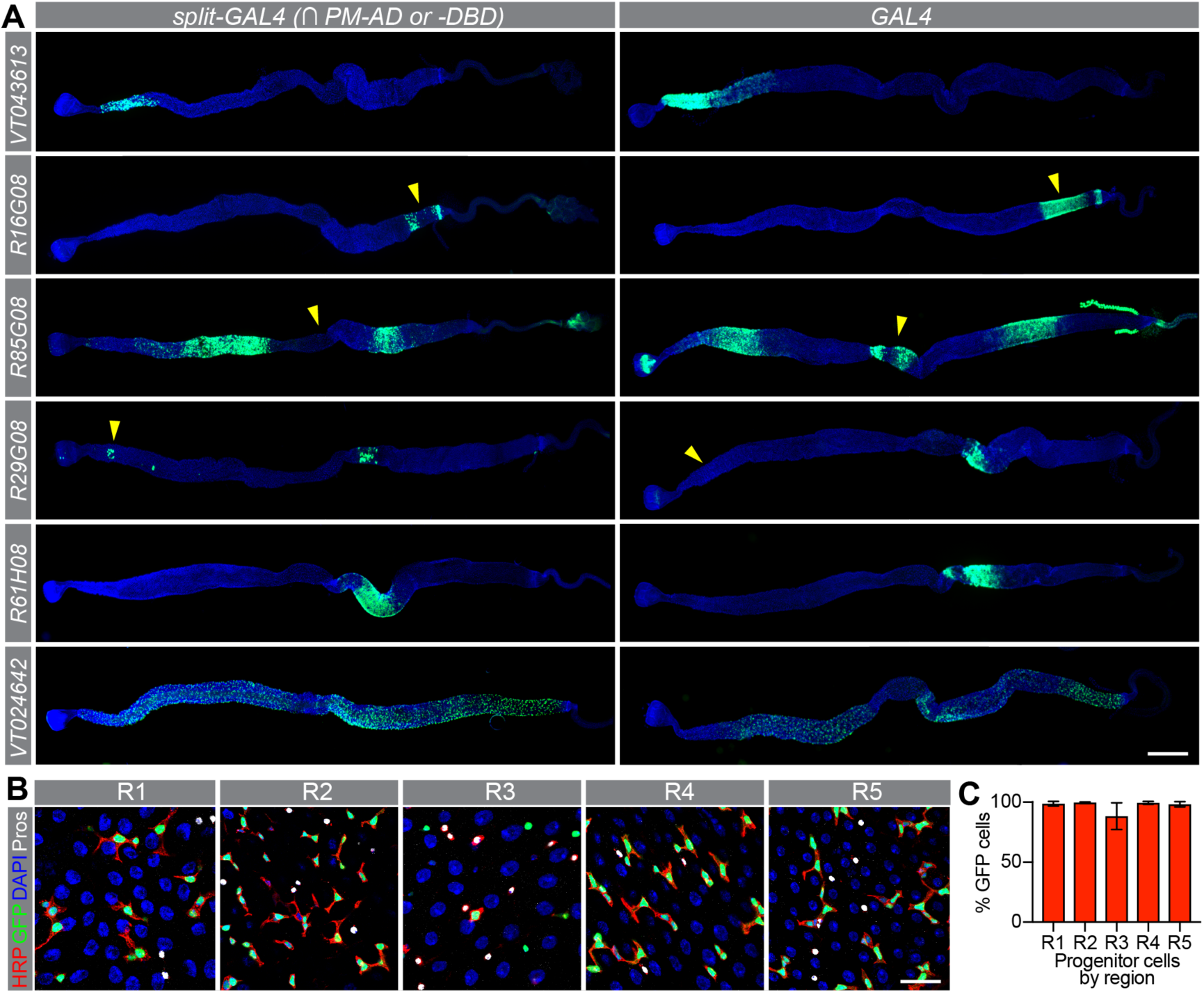
Enhancer fragments can confer similar expression patterns in different transgenic contexts. (A) *UAS-Stinger* expression (green) in adult female intestines when driven by six enhancer fragments in the split-GAL4 (left) or GAL4 (right) systems. Yellow arrowheads indicate minor expression differences. DAPI staining is in blue. (B) *UAS-Stinger* expression (green) driven by *VT024642-GAL4* in intestinal regions R1–5 with counterstaining for progenitor cells (anti-HRP in red), EEs (anti-Pros in white), and all cell nuclei (DAPI in blue). (C) Quantification of progenitor cell labeling by *VT024642-GAL4* driving *UAS-Stinger* expression in R1–R5. Scale bars; 500μm (A), 25μm (B). Complete genotypes are listed in Table S2.

## DISCUSSION

Here, we describe the expression of 424 split-GAL4 drivers in the Drosophila adult midgut that, altogether, represent 352 distinct fragments of regulatory sequence lying within 10 kb of 907 genes. By pairing these drivers with reference drivers specific to intestinal cell types, we describe 653 split-GAL4 combinations that identify a myriad of region-, cell type-, and sex-specific cell populations. Descriptions of these patterns are available both in File S2 and online at https://bdsc.indiana.edu/stocks/gal4/midgut_splitgal4.html, allowing the design of additional driver pairings to target particular intestinal cells, as well as the creation of new, cell-specific transgenes incorporating the enhancer sequences.

This new resource will provide tools for manipulating specific cell subtypes with more precision. We identified *R10F08*-*p65AD* as an example of a driver expressed in a novel cell population and we anticipate that split-GAL4 drivers will also identify subsets of cells based on physiological state, such as age, disease, or response to environmental conditions. Integrating split-GAL4 drivers into experiments with sophisticated molecular assays seems particularly promising. For example, one or more of the 83 region-specific, EC-specific split-GAL4 combinations described here might specifically label some of the regional EC subtypes recently identified using single-cell RNA sequencing (scRNA-seq) analysis (Hung et al., 2020). Indeed, we identified six enhancer fragments from drivers expressed in region-specific EC subpopulations that originated within 10 kb of genes expressed in single EC subtypes in the scRNA-seq analysis (Table 4), and several more fragments associated with genes expressed in a limited number of subtypes. While the relationships of driver expression, EC subtype and specific genes need to be confirmed, these results show that split-GAL4 combinations have tremendous potential for experiments exploring the association of transcription factors and enhancers in specific intestinal cell populations.

**Table 4.** Drivers with enhancer fragments associated with EC subtype-specific genes

| Region-specific EC driver | EC subtype-specific gene | EC subtype |
| --- | --- | --- |
| *P{VT019063-GAL4.DBD}attP2* | *5-HT1A* | pEC2 |
| *P{R61H08-GAL4.DBD}attP2* | *AstCC* | LFC |
| *P{VT063961-GAL4.DBD}attP2* | *CCKLR-17D3* | LFC |
| *P{VT020029-GAL4.DBD}attP2* | *hbn* | pEC1 |
| *P{VT017655-GAL4.DBD}attP2* | *Sema2b* | pEC3 |
| *P{R31E11-p65.AD}attP40* | *Ubx* | aEC2 |

This work expands the number of defined enhancer sequences that label entire sets of intestinal cell types. Prior analyses had identified enhancer sequences from *miranda* and *zfh2* that label progenitor cells (Bardin et al., 2010; Rojas Villa et al., 2019), and here we identified two more enhancer-containing fragments labeling progenitor cells: *VT024642* from the region of *Acp62F* and *VT004241* from the region of *Akap200*. All four of these chromosomal segments contain binding sites for the *dorsal* transcription factor (ModEncode Consortium et al., 2010), suggesting a regulatory logic that could be explored. We also identify *VT004958* as a genomic fragment containing enhancer sequences active and specific to all ECs. In contrast to progenitor and EC cells, however, we failed to identify regulatory sequences active in all EEs—despite testing split-GAL4 and GAL4 drivers with six different genomic fragments associated with the EE-specific *pros* gene. Testing additional enhancer fragments and fragment combinations from *pros* may eventually identify a pan-EE element. Finally, we did not identify a regulatory fragment specific to either ISCs or EBs, consistent with recent sequencing results indicating that the transcriptomes of ISCs and EBs are highly similar (Dutta et al., 2015; Hung et al., 2020).

We also made observations that highlight the complexities of the split-GAL4 system. For example, 47 split-GAL4 drivers displayed expression with the PM reference drivers in the primary screen but did not promote expression with any of the cell type reference drivers in the secondary screen. In addition, we noted some drivers expressed in different patterns with the P, EE, and EC reference drivers than we expected from their expression patterns with a PM reference driver. Finally, we saw that split-GAL4 patterns can differ depending on the choice of *UAS-responder*. Altogether, these observations indicate that additional variables—such as genetic background, relative enhancer strength, and insertion site—may impact split-GAL4 transgene expression and should be considered when using the resources reported here.

We expect that this resource will have broad utility in experiments to label and/or, genetically manipulate distinct subsets of intestinal cells. The split-GAL4 drivers used here have the disadvantage of not being compatible with popular gene expression systems employing GAL80 repression, because the drivers utilize the p65 activation domain, which, unlike the GAL4 activation domain, does not bind GAL80 (Dionne et al., 2018; Ma and Ptashne, 1987). Nevertheless, as we have shown, enhancer fragments identified using the split-GAL4 system can be transferred to transgenic constructs compatible with GAL80 and to other gene expression systems for fine-tuned conditional or intersectional control. This resource can be expanded still by constructing additional split-GAL4 drivers, including ones that contain regulatory elements from genes known to be expressed in distinctive patterns in the intestine (Buchon et al., 2013; Dutta et al., 2015; Hung et al., 2020; Marianes and Spradling, 2013). It should also prove valuable in future experiments combining the split-GAL4 system with other gene expression systems to target multiple cell types simultaneously. Such efforts will continue to enhance the capability of Drosophila experimental systems for studying conserved core biological processes.

## ACKNOWLEDGEMENTS

We thank Steve Stowers, the Developmental Studies Hybridoma Bank, and the Drosophila Genome Resource Center (supported by NIH grant P40OD010949) for reagents; the Light Microscopy Imaging Center (supported by NIH grant S10OD024988) for access to the SP8 confocal; Lawrence Washington for preliminary studies; Annette Parks and Jim Smotherman for help constructing the accompanying webpages; our colleagues at Indiana University for helpful discussions; and the National Institutes of Health for financial support (awards R21OD026525 to KRC and NSS, R01GM124220 to NSS, and P40OD018537 to KRC). The authors declare no competing financial interests.

---

**Figure S1.**
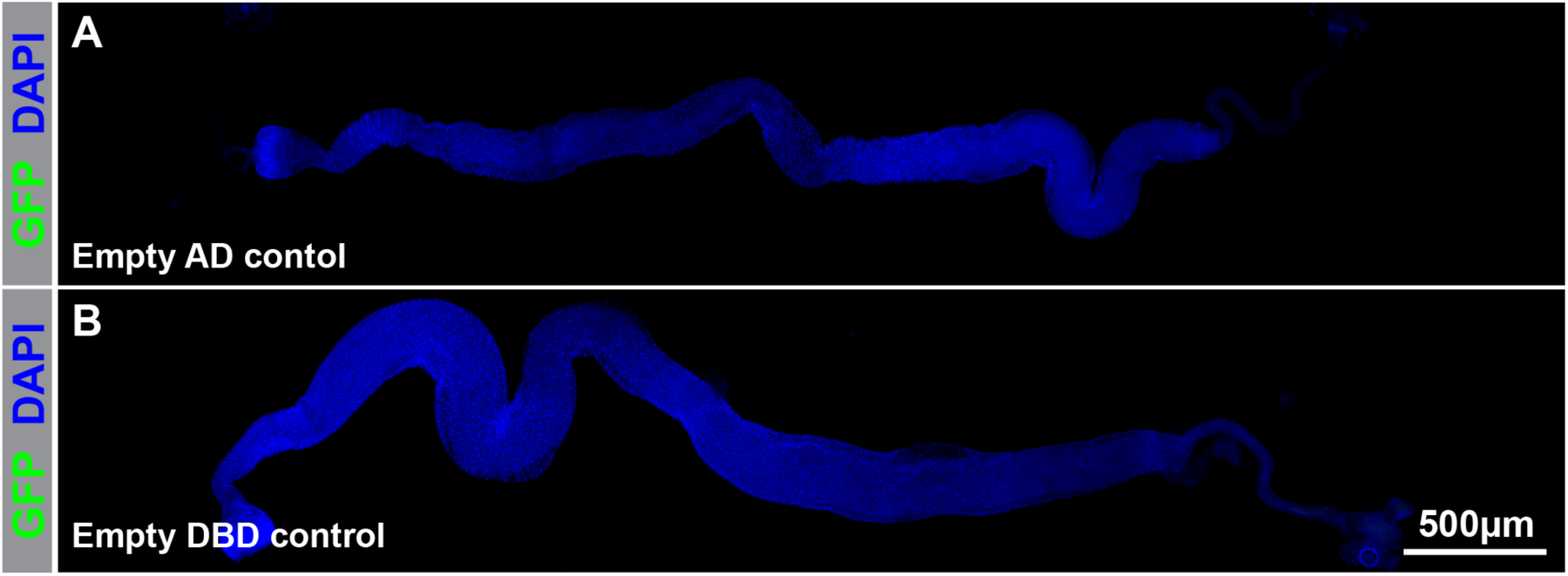
GFP expression in the presence of pan-midgut split-GAL4 drivers has little background. (A, B) Lack of *UAS-6XGFP* intestinal expression in the presence of control drivers lacking enhancer fragments (*CG10116-GAL4DBD* ∩ *empty-p65AD* in A and *CG10116-p65AD* ∩ *empty-GAL4DBD* in B). Complete genotypes are listed in Table S2.

**Figure S2.**
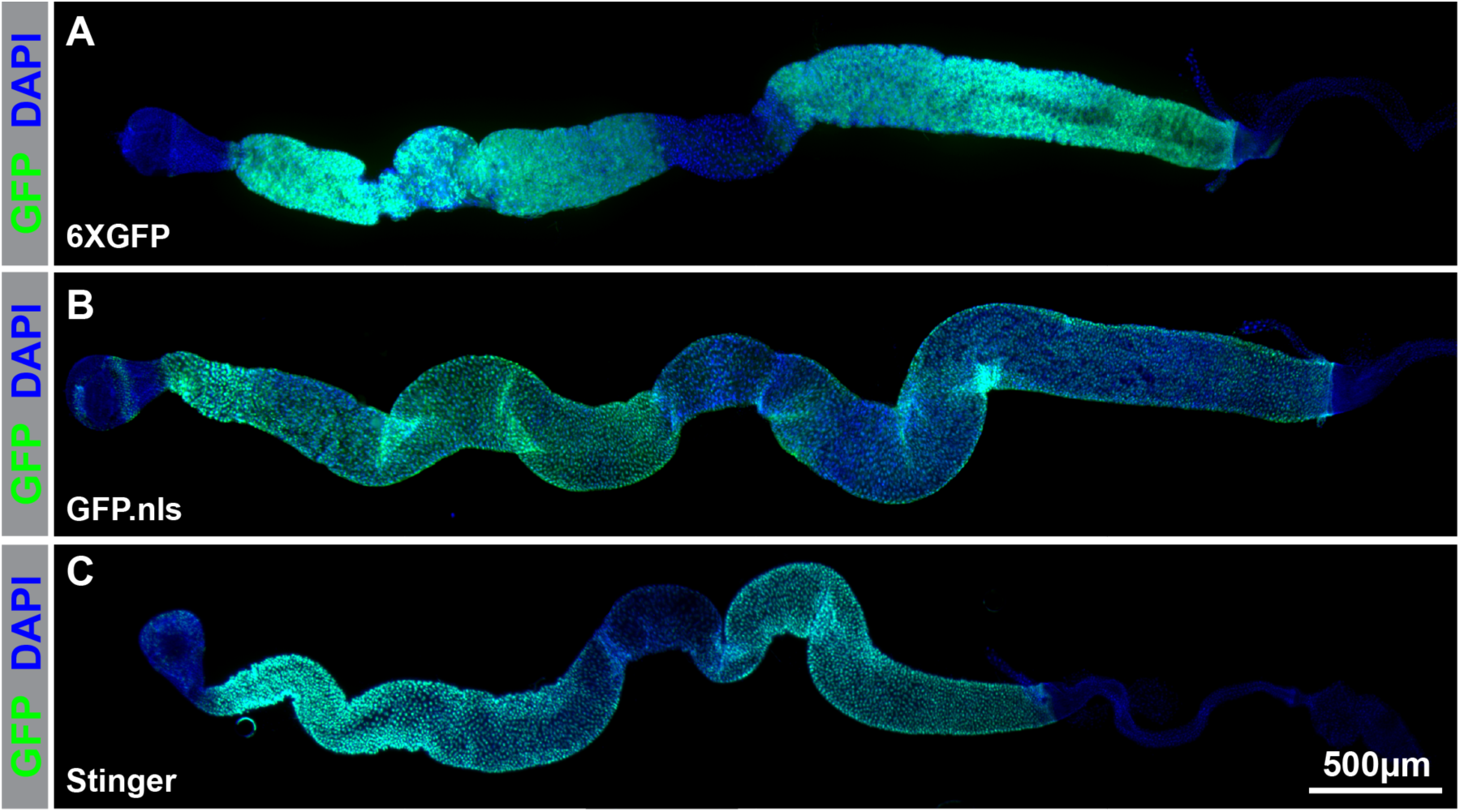
GFP expression varies by *UAS-GFP* transgene. (A–C) GFP expression in the presence of pan-midgut split-GAL4 drivers (*CG10116-p65AD* ∩ *CG10116-GAL4DBD*) and *P{20XUAS-6XGFP}* (A), *UAS-GFP.nls* (B), or *UAS-Stinger* (C). Sample in B was stained with anti-GFP antibody while samples in other panels were not. Note that GFP expression in R3 is much higher with *UAS-GFP.nls* (B) compared to the other responder transgenes (A, C). Complete genotypes are listed in Table S2.

**Figure S3.**
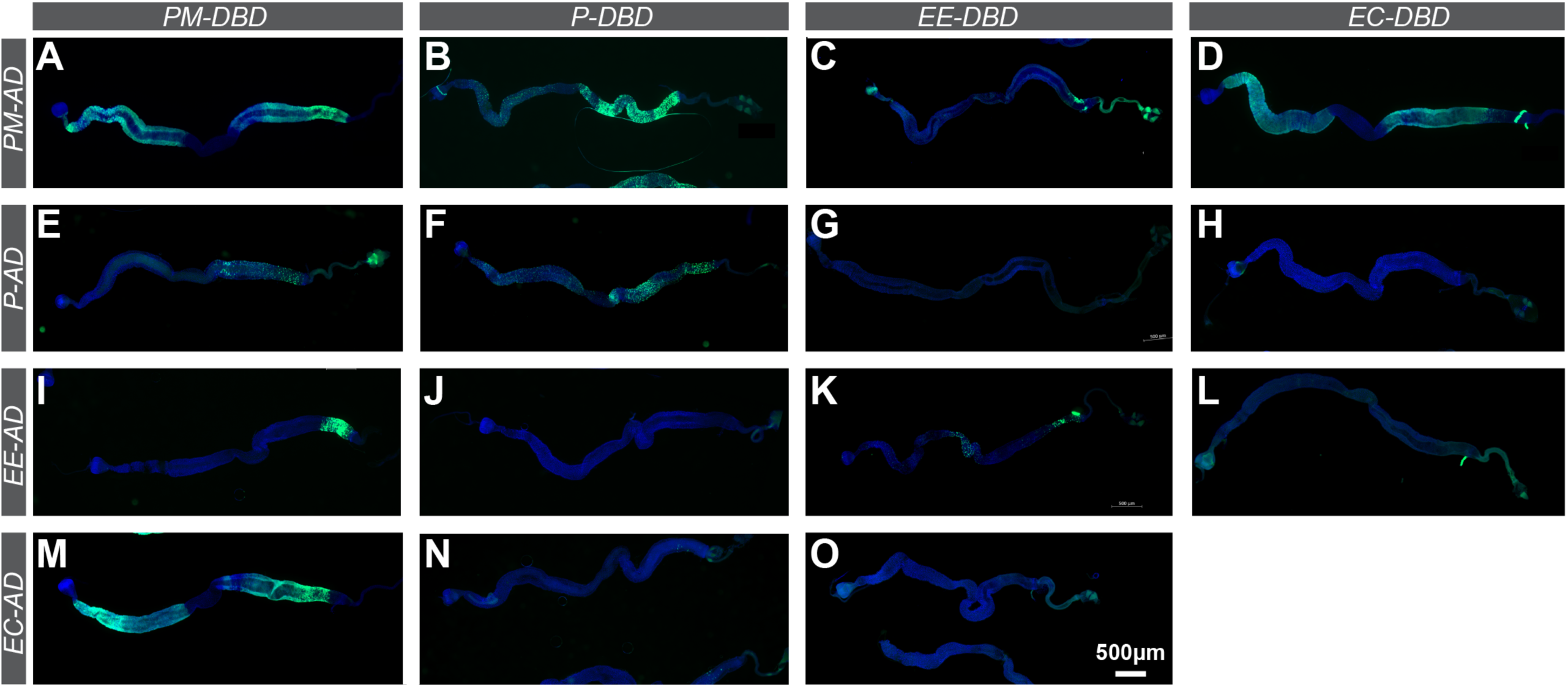
GFP expression in the presence of reference driver combinations and *UAS-6XGFP*. (A–O) In general, combinations of drivers expected to express in different cell types failed to drive expression of *UAS-6XGFP* (green) while drivers expected to express in the same cell types expressed *UAS-6XGFP*. PM drivers promoted *UAS-6XGFP* expression in R3 poorly (A–D, E, I, M). Large cells in R4 and R5 occasionally expressed *UAS-6XGFP* unexpectedly in driver combinations involving P and EE drivers (N and L). Intestines counterstained with DAPI (blue). Complete genotypes are listed in Table S2.

